# XBP1s-Mediated ER Proteostasis Network Enhancement Can Selectively Improve the Folding and Secretion of an Osteogenesis Imperfecta-Causing Collagen-I Variant

**DOI:** 10.1101/2021.04.15.439909

**Authors:** Andrew S. DiChiara, Ngoc-Duc Doan, Agata A. Bikovtseva, Lynn Rowley, Vincent L. Butty, MaryAnn E. Weis, David R. Eyre, Shireen R. Lamandé, John F. Bateman, Matthew D. Shoulders

## Abstract

Osteogenesis imperfecta (OI) is typically caused by autosomal dominant mutations in genes encoding collagen type-I, most commonly resulting in Gly→Ser triple-helical domain substitutions that disrupt collagen folding and/or stability. Here, we test the hypothesis that upregulating the endoplasmic reticulum (ER) proteostasis network via the unfolded protein response (UPR) can improve the folding and secretion of the clinically severe, prototypical OI-causing *COL1A1* p.G425S collagen-α1(I) variant. We first show that small molecules that activate the entire UPR by causing global ER protein misfolding stress severely ablate collagen-I secretion from both G425S Colα1(I)- and wild-type (WT) Colα1(I)-expressing primary fibroblasts. In contrast, stress-independent, specific induction of just the UPR’s XBP1s transcriptional response can enhance collagen-I secretion from G425S Colα1(I) patient primary fibroblasts up to ~300% of basal levels. Notably, the effect is selective – collagen-I secretion from WT Colα1(I)-expressing healthy donor primary fibroblasts is unaltered by XBP1s. XBP1s pathway activation appears to post-translationally enhance the folding/assembly and secretion of G425S Colα1(I), as only modest impacts on collagen-I transcription or synthesis are observed. Consistent with this notion, we find that the stable, triple-helical collagen-I secreted by XBP1s-activated G425S α1(I) patient fibroblasts includes a higher proportion of the mutant α1(I) polypeptide than the collagen-I secreted under basal ER proteostasis conditions. We note that consistent reproducibility of these results is dependent on as yet unascertained experimental variables. Still, these promising observations suggest the potential for ER proteostasis network modulation to improve mutant collagen proteostasis in the collagenopathies, motivating further investigation of the effect’s generality, underlying mechanism, and potential therapeutic benefits.

## INTRODUCTION

Mutations that result in dysregulated collagen proteostasis, defined as an inability of cells to handle and produce appropriate quantities of folded and functional collagen, cause an array of diseases known as the collagenopathies^1, 2^. The collagenopathies have divergent clinical phenotypes, ranging from mild to severe, depending on the specific mutation and the type of collagen involved. The prototypical collagenopathy, osteogenesis imperfecta (OI), is characterized by symptoms such as bone fragility, reduced bone mass, and short stature^3^. As is also true for the other collagenopathies, current OI therapies remain inadequate for alleviating both skeletal and non-skeletal manifestations of the disease.

OI is most often caused by autosomal dominant mutations in the *COL1A1* or *COL1A2* genes that encode the procollagen-α1(I) and procollagen-α2(I) strands of the 2:1 proα1(I):proα2(I) heterotrimeric procollagen molecule (procollagen-I refers to collagen-I prior to the extracellular cleavage of its N- and C-propeptides)^3^. The most common class of OI-causing mutations are missense mutations causing substitution of various conserved Gly residues within the central, functionally critical triple-helical domain of collagen-I^4^. A Gly residue is required at every third position within the triple-helical domain of mature collagen polypeptides, because it is the only amino acid small enough to be accommodated in the interior of the sterically occluded triple helix^5^. Substitution of Gly with any other amino acid creates a steric “bump” in the folded triple helix, disrupts the stabilizing ladder of interstrand N–H_(Gly)_…O=C_(Xaa)_ hydrogen bonds, and delays triple-helix folding (**Figure 1**)^5, 6^.

**Figure 1.**
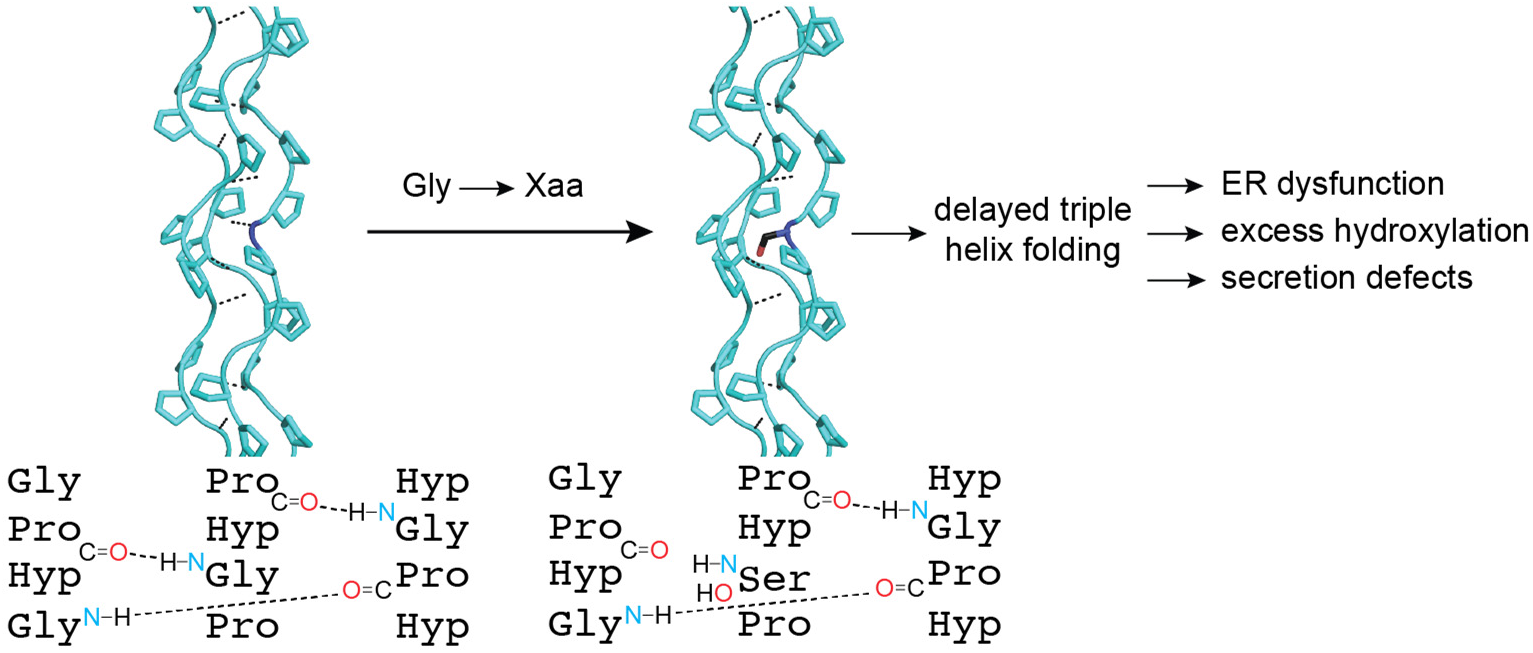
High-resolution crystal structure of a collagen triple helix formed from (Pro-Pro-Gly)_10_ (PDB1K6F)^87^. An exemplary glycine to serine substitution and resulting hydrogen bond disruption is shown, obtained by replacing a single Gly residue with Ser. A diagram of the staggered arrangement of the three distinctive polypeptide strands in a segment of a triple helix and the ladder of interstrand hydrogen bonds that is disrupted by a Gly→Ser substitution is also shown.

OI-causing mutations can engender at least three defects that pathologically disrupt collagen proteostasis: (1) Non-functional collagen may be allowed to escape the cell instead of being subjected to quality control. Secreted, non-functional collagens may cause bone defects^7–9^ or disrupt essential extracellular matrix protein– protein and/or protein–cell interactions^4, 10, 11^. (2) Collagen-I mutations could result in insufficient production of functional collagen-I, owing to hyperactive quality control and inefficient folding. A reduction in functional collagen-I production itself can be damaging^12, 13^. Slow collagen-I folding and secretion often also results in over-hydroxylation of triple-helical lysine residues^14–16^, which may result in excessive collagen-I crosslinking, hyper-*O*-glycosylation, and consequent pathologic structural defects in bone and other tissues^17^. (3) Misfolding or slow-folding collagen-I could overwhelm the endoplasmic reticulum (ER) proteostasis network, resulting in intracellular collagen accumulation and chronic cell stress, likely promoting the apoptosis of collagen-I producing osteoblasts and thereby reducing bone health^18–21^. The resultant ER dysfunction could also lead to a net decrease in collagen-I production that exceeds the consequences of haploinsufficiency mutations in collagen-I (e.g., premature stop codons). Haploinsufficiency mutations reduce the net synthesis of collagen-I but do not create a folding challenge for the ER and cause only mild OI^3, 12^.

Distinctive OI mutations likely lead to disease by various combinations of one or more of these mechanisms. The integral role of dysregulated collagen proteostasis in all these mechanisms highlights the need to develop strategies that can alleviate such defects, if possible^1, 2, 19^. The endogenous cellular signaling pathway typically responsible for addressing defects in ER proteostasis is the unfolded protein response (UPR)^22, 23^. The UPR consists of three distinct arms, ATF6, IRE1-XBP1s, and PERK-ATF4, that cooperate to resolve ER proteostasis defects by transcriptionally remodeling the ER’s chaperone and quality control network while simultaneously reducing translation to lower the client protein load on the ER. The predominant function of the ATF6 and IRE1-XBP1s UPR arms is to restore proteostasis by transcriptionally upregulating ER chaperones and quality control machinery through the activity of the cleaved ATF6(1–373) and XBP1s transcription factors. The PERK UPR arm regulates ER client protein translation and, if chronically activated, ultimately causes cellular apoptosis.

Misfolding- or aggregation-causing mutations in ER proteins can, at least in some cases, lead directly to UPR induction^24^. The resulting UPR activation, by reshaping the ER proteostasis network to better support protein folding and production, can prove beneficial to help address the underlying proteostasis defect, although long-term chronic ER stress is problematic^23, 25^. Regardless, UPR activation by the common disease-causing, triple-helical Gly substitutions in collagen-α1(I) and collagen-α2(I) (α1(I) and α2(I), respectively) is not generally observed^18, 19, 26–28^. The lack of UPR activation despite the presence of a large quantity of a misfolding client protein may reflect an inability of the ER’s protein misfolding sensors to detect accumulation of an atypical misfolded fibrous protein, rather than a globular protein with a hydrophobic core.

The absence of a prototypical UPR in most collagenopathies could present an opportunity for therapeutic intervention. Specifically, there is still the possibility that enhancing the ER proteostasis network via artificial UPR activation could prove beneficial for resolving collagen-I proteostasis defects, for example by improving folding and trafficking of OI variants. Indeed, in a number of protein misfolding-related disease contexts, genetic or pharmacologic UPR modulation has been shown to address underlying proteostasis defects^29–34^. For example, the UPR-mediated ER proteostasis network upregulation induced by treatment with chemical ER stressors enhances the folding and trafficking of misfolding-prone lysosomal enzyme variants to the lysosome, where they are sufficiently active to rescue lysosome function^35, 36^. This enhanced trafficking also relieves ER dysfunction caused by the poorly trafficked enzyme variants.

Here, we show that remodeling the ER proteostasis network via forced expression of the UPR’s XBP1s transcription factor can, under some conditions, improve the folding and secretion of collagen-I by OI patient primary fibroblasts. When an increase in collagen-I secretion is detectable, we observed that XBP1s activation is selectively enhancing mutant collagen-I proteostasis, consistently having no significant impact on the folding and trafficking of collagen-I by wild-type (WT) α1(I)-expressing primary fibroblasts. The effect appears to be due to a post-translational enhancement of mutant collagen-I folding, assembly, and trafficking, as XBP1s only minimally influences collagen-I transcription or translation. Consistent with this observation, we observed that the enhanced secretion of collagen-I molecules from XBP1s-treated primary OI patient fibroblasts results in increased incorporation of G425S α1(I) into secreted, stable triple helices. These observations provide support for the notion that pharmacologic targeting of the ER proteostasis network can address disease-causing collagen proteostasis defects.

## MATERIALS AND METHODS

### Cell Culture

WT α1(I)-expressing (GM05294; healthy donor) or G425S (numbering from the first residue of the N-propeptide; this mutation corresponds to G247S when numbering is started at the first glycine of the triple helix) α1(I)-expressing (GM05747; OI patient) primary fibroblasts were obtained from Coriell Cell Repository and cultured in complete MEM (Corning) supplemented with 15% fetal bovine serum (FBS; Corning), 100 IU penicillin (Corning), 100 μg/mL streptomycin (Corning), and 2 mM L-glutamine (Corning). For sub-culturing, cells were trypsinized and resuspended in ample fresh media, collected by centrifugation, and resuspended in complete fresh media to count and then plate cells for experiments. Cells were counted using the Countess II FL (Life Technologies). Collagen-I production was induced using 200 μM sodium ascorbate (Amresco). Cell viability assessment was performed using the CellTiter-Glo® assay (Promega) according to the manufacturer’s instructions.

### Cell Lysis, SDS-PAGE, and Immunoblotting

After trypsinization and neutralization in complete media, intact cells were collected by centrifugation, washed with 1× PBS, and then lysed in radio-immunoprecipitation assay buffer (RIPA) containing 150 mM sodium chloride, 50 mM Tris-HCl at pH 7.5, 1% Triton-X 100 (Integra), 0.5% sodium dodecyl sulfate, 0.1% sodium deoxycholate, protease inhibitor tablets (Pierce), and 1 mM phenylmethylsulfonyl fluoride. Cells were lysed on ice for 20 min, the lysate was cleared by centrifugation at 16,800 rpm at 4 °C, and then protein concentration in the supernatant was quantified and normalized prior to separation under reducing conditions on 8% SDS-PAGE. Identical volumes of media were also collected and used directly for SDS-PAGE. For immunoblotting, proteins were transferred to nitrocellulose membranes using the TransBlot Turbo system (Bio-Rad) using transfer buffer at pH 9.2 containing 48 mM Tris, 39 mM glycine, 0.1% SDS, and 20% ethanol. Membranes were blocked in 5% milk and then probed with the indicated primary antibodies diluted in 5% bovine serum albumin (VWR). Blots were imaged after incubation with appropriate primary and secondary antibodies by scanning on an Odyssey IR imager (LI-COR). Antibodies used were: rabbit anti-α1(I) [LF68] (Kerafast ENH018-FP) raised against the C-telopeptide of the α1(I) chain (Figures: **2a**, **3**, and **S2**); rabbit anti-Colα1(I) (MyBioSource MBS502153) raised against the C-telopeptide of the α1(I) chain (Figure: **2c**); rabbit anti-α2(I) (Sigma SAB4500363) raised against human α2(I) triple-helical domain-derived peptides (Figures: **2a**, **3**, **5a**, and **S2**). Secondary antibodies were obtained from LiCor Biosciences: 800CW goat anti-rabbit, 800CW goat anti-mouse, 680LT goat anti-rabbit, 680LT goat anti-mouse.

**Figure 2.**
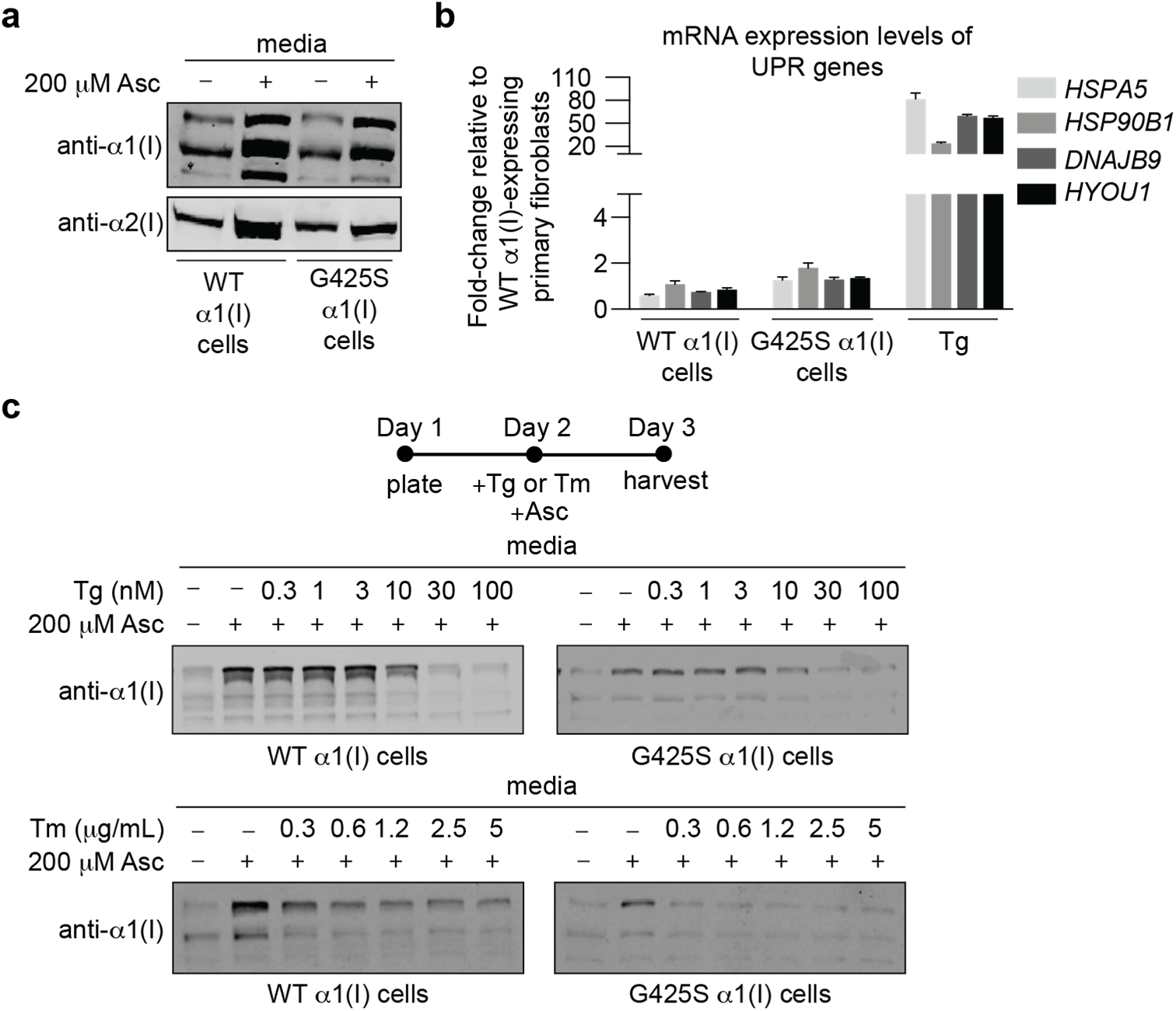
**a.** Immunoblot showing secretion of α1(I) and α2(I) from WT α1(I) and G425S α1(I) primary patient fibroblasts. Equal numbers of cells were plated and collagen-I production was initiated by treatment with sodium ascorbate (Asc) for 24 h. The presence of multiple α1(I) bands corresponds to proα1(I) and α1(I) before and after cleavage of the C-propeptide and/or the N-propeptide. **b.** qPCR analysis of unfolded protein response (UPR)-regulated transcripts in WT α1(I) and G425S α1(I) primary fibroblasts. Fold-change in transcript levels was normalized to WT α1(I) fibroblasts not treated with sodium ascorbate. Thapsigargin (Tg) treatment was used as a positive control for UPR activation. Error bars represent SD across three technical replicates. **c.** Immunoblot showing secretion of α1(I) from WT α1(I) and G425S α1(I) primary fibroblasts treated with increasing concentrations of Tg (**top**) or tunicamycin (Tm) (**bottom**). One day post-plating, cells were treated with Tg or Tm and α1(I) secretion was initiated by treatment with Asc, followed by harvesting for analysis after 24 h.

### Adenovirus Transductions

The ViraPower^TM^ Adenoviral Gateway^TM^ system was used to generate replication-incompetent adenoviruses encoding either XBP1s, eGFP, DHFR.ATF6(1–373),^31^ or DHFR.YFP. Viruses were titered using the Adeno-X qPCR Titration kit (Clontech), according to the manufacturer’s instructions. WT α1(I)- or G425S α1(I)-expressing primary fibroblasts (1.0 × 10^5^ cells) were plated the day before transduction in a 6-well plate. 24 h post-plating, media was replaced and cells were treated with the indicated adenovirus at an experimentally optimized multiplicity of infection determined for each batch of adenovirus used. Following 24 h of incubation with virus, media was replaced. 48 h post-media change, the media was replaced with fresh media supplemented with 200 μM Asc. Following ~24 h of incubation with Asc, cells were harvested by trypsinization. For experiments involving DHFR.ATF6(1–373) or DHFR.YFP, WT Colα1(I)-expressing or G425S α1(I)-expressing primary fibroblasts (2.0 × 10^5^ cells) were plated the day before transduction in a 6-well plate. 24 h post-plating, the media was replaced and supplemented with DHFR.ATF6(1–373) or DHFR.YFP adenoviral supernatant at an experimentally optimized MOI yielding TMP-dependent activation of the ATF6 arm of the UPR. Following 24 h of incubation with virus, cells were split and then supplemented with media containing 10 μM TMP or vehicle. 48 h post-media change or split, media was removed and replaced with 1 mL of fresh media supplemented with 200 μM ascorbate. Media and cells were harvested for downstream analysis after ~18 h.

### Quantitative RT-PCR

Relative mRNA expression levels of genes of interest were assessed by quantitative RT-PCR. Cells were harvested by trypsinization and washed with PBS. Total RNA was extracted using the Omega RNA Purification Kit. RNA concentrations were quantified and normalized to 1 μg total RNA for cDNA reverse transcription. cDNA was synthesized in a BioRad Thermocycler using the Applied Biosystems reverse transcriptase cDNA kit. KAPA BioSystems Sybr Fast qPCR master mix, appropriate primers (Sigma Aldrich; see **Table S3**), and cDNA were then used for amplification in a Light Cycler 480 II real-time PCR Instrument. Primer integrity was assessed by thermal melt to ensure homogeneity. Transcripts were normalized to the housekeeping gene RPLP2. Standard deviation for *n* = 3 is shown in plots.

### RNA-Seq

Cells were transduced with Ad.GFP or Ad.XBP1s and then treated with Asc as described above. Following treatment with Asc, cells were harvested by trypsinization, washed with PBS, and total RNA was extracted using the Omega RNA Purification Kit. RNA samples were quantified and quality assessed using an Advanced Analytical Fragment Analyzer. Initial steps in library preparation were then performed on a Tecan EVO150. 10 ng of total RNA was used for library preparation. 3′-DGE-custom primers 3V6NEXT-bmc#1–12 were added to a final concentration of 1 μM (5’-/5Biosg/ACACTCTTTCCCTACACGACGCTCTTCCGATCT[BC_6_]N_10_T_30_VN-3’ where 5Biosg = 5’-biotin, [BC6] = 6 bp barcode specific to each sample/well, N10 = Unique Molecular Identifiers, Integrated DNA Technologies). After addition of the oligonucleotides, Maxima H Minus RT was added per the manufacturer’s recommendations with the template-switching oligo 5V6NEXT (10 μM, [5V6NEXT: 5’-iCiGiCACACTCTTTCCCTACACGACGCrGrGrG-3’ where iC: iso-dC, iG: iso-dG, rG: RNA G]) followed by incubation at 42 °C for 90 min and inactivation at 80 °C for 10 min. Following the template-switching reaction, cDNA from 12 wells containing unique well identifiers were pooled together and cleaned using RNA Ampure beads at 1.0×. cDNA was eluted with 17 μL of water followed by digestion with exonuclease I at 37 °C for 30 min, and then inactivated at 80 °C for 20 min. Second strand synthesis and PCR amplification was performed by adding the Advantage 2 Polymerase Mix (Clontech) and the SINGV6 primer (10 pmol, Integrated DNA Technologies 5’-/5Biosg/ACACTCTTTCCCTACACGACGC-3’) directly to the exonuclease reaction. 15 cycles of PCR were performed followed by clean-up using regular SPRI beads at 0.6×, and eluted with 20 μL of elution buffer. Successful amplification of cDNA was confirmed using a Fragment Analyzer. Illumina libraries were then produced using standard Nextera tagmentation, substituting the P5NEXTPT5-bmc primer (25 μM, Integrated DNA Technologies (5’-AATGATACGGCGACCACCGAGATCTACACTCTTTCCCTACACGACGCTCTTCCG*A*T*C*T*-3’ where * = phosphorothioate bonds) in place of the normal N500 primer. Final libraries were cleaned using SPRI beads at 0.7× and quantified using a Fragment Analyzer and qPCR before being loaded for paired-end sequencing using the Illumina NextSeq500 in paired-end mode (20/50 nt reads).

Post-sequencing, quality-control on each of the libraries was performed to assess coverage depth, enrichment for messenger RNA (exon/intron and exon/intergenic density ratios), fraction of rRNA reads and number of detected genes using bespoke scripts. Read pairs were combined into a single Fastq with well/unique molecular identifier (UMI) information concatenated with the second read name. Reads sharing sequence and UMI information were collapsed into single exemplars. Sequences were aligned against the human genome GRCh38/hg38 ENSEMBL 89 using STAR ^37^, with parameters --runThreadN 8 --runMode alignReads --outFilterType BySJout --outFilterMultimapNmax 20 --alignSJoverhangMin 8 --alignSJDBoverhangMin 1 --outFilterMismatchNmax 999 --alignIntronMin 10 --alignIntronMax 1000000 --alignMatesGapMax 1000000 --outSAMtype BAM SortedByCoordinate --quantMode TranscriptomeSAM, pointing to a Suffix Array assembly containing exon-exon junctions for 75 nucleotide reads. Gene expression was estimated based on reads mapping near the 3’ end of transcripts using ESAT ^38^ with the following parameters -task score3p -alignments $sample_list -wLen 50 -wExt 5000 -wOlap 0 -sigTest 0.01 -multimap ignore, pointing to a refseq-based gene annotation (hg38, as downloaded from the UCSC Genome Browser) and gene-level data were used for downstream analyses.

Differential expression was performed in the R statistical environment (R v. 3.4.0) using Bioconductor’s DESeq2 package on all annotated genes^39^. Dataset parameters were estimated using the estimateSizeFactors(), and estimateDispersions() functions. Read counts across conditions were modeled based on a negative binomial distribution and a Wald test was used to test for differential expression (nbinomWaldtest(), all packaged into the DESeq() function), using the treatment type as a contrast and the replicate of origin as a factor. Fold-changes, *p*-values and Benjamin-Hochberg-adjusted *p*-values (BH) are reported for each gene. Regularized fold-changes were calculated using the lfcShrink() function in “normal” mode.

GSEA was performed to compare genes upregulated upon XBP1s induction in WT vs G425S α1(I) primary fibroblast cells. Differential expression results from DESeq2 were retrieved, and the “stat” column was used to pre-rank genes for GSEA. Briefly, these “stat” values reflect the Wald’s test performed on read counts as modeled by DESeq2 using the negative binomial distribution. Genes that were not expressed were excluded from the analysis. GSEA (desktop version, v3.0)^40, 41^ was run in the pre-ranked mode against MsigDB 7.0 C5 (gene ontology) sets, using the official gene symbol as the key, with a weighted scoring scheme, normalizing by meandiv, with gene sets between 10 and 2000 genes (9438 gene sets retained for C5). 5000 permutations were run for *p*-value estimation. GSEA enrichment plots were replotted using a modified version of the ReplotGSEA.R script.

### Metabolic Labeling

72 h post-transduction with Ad.GFP or Ad.XBP1s adenoviruses, media was removed from WT α1(I) or G425S α1(I) primary fibroblasts and replaced with fresh media containing 200 μM ascorbate. 2 h later, media was replaced with fresh media containing 200 μM ascorbate and 100 mCi/mL of ^35^S-Cys/Met (Perkin Elmer) for 10 min. Following labeling, media was removed, cells were washed 2× with PBS, and then lysed immediately in the plate using RIPA. Following 20 min of lysis on ice, samples were collected and cell debris was removed by centrifugation at 10,000 x *g* and 4 °C for 15 min. Supernatants were collected, split equally into two tubes, and supplemented with 8× collagenase buffer containing 50 mM HEPES at pH 7.5 and 360 μM calcium chloride. Half of the samples were treated with a final concentration of 5 μg/mL bacterial collagenase (Worthington Biochemical) for 3 d at 37 °C. Samples were then loaded onto homemade 10% SDS-PAGE gels. Gels were dried, developed for 2 d, and imaged using a phosphor screen and a Typhoon 7800 imager. Band intensities were quantified using ImageQuant TL (GE Healthcare). Experiments were performed in biological triplicate with standard deviation shown.

### Secreted Collagen-I Assessment

WT α1(I)-expressing or G425S α1(I)-expressing primary fibroblasts were plated in a 6-cm dish, and then transduced 24 h later with Ad.GFP or Ad.XBP1s. 24 h post-transduction, media was changed. 72 h post-transduction, collagen production was induced by treatment with 200 μM ascorbate for 24 h. Media was collected, chilled at 4 °C, and treated with 1 mM PMSF, 176 mg/mL ammonium sulfate, and 100 mM Tris-HCl at pH 7.4 to precipitate the secreted procollagen-I. Samples were mixed end over end overnight at 4 **°**C, and pelleted by centrifugation at 10,000 x *g* and 4 °C for 30 min. Supernatant was removed and the remaining pellet was resolubilized in 400 mM NaCl and 100 mM Tris-HCl at pH 7.4. Collagen-I samples were then aliquoted and treated with 0.1 mg/mL trypsin or 0.2 mg/mL chymotrypsin (Sigma Aldrich)^42^. Samples were analyzed by SDS-PAGE using an antibody raised against peptides derived from the triple-helical domain of α2(I).

### Mass Spectrometry Analysis of Secreted G425S Colα1(I)

G425S α1(I) primary patient fibroblasts (1.0 × 10^5^ cells) were plated in a 6-well plate and transduced 24 h later with Ad.GFP or Ad.XBP1s. 24 h post-transduction, media was changed. 72 h post-transduction, collagen production was initiated by treating with 200 μM ascorbate for 24 h. Media samples were then collected, chilled at 4 °C, and treated with 1 mM PMSF. Samples were acidified to 3% acetic acid and treated with pepsin (~0.1 mg/ml and ~1:10 pepsin:collagen) for 24 h^43^. Collagen was then precipitated by addition of sodium chloride to a 0.7–0.9 M final concentration. The resulting collagen was separated on a homemade 6% SDS-PAGE gel. The gel was stained with Coomassie dye and the α1(I) band was excised, reduced with DTT, alkylated with iodoacetamide, and trypsinized overnight. The resultant tryptic peptides were analyzed by LC-MS/MS analysis to determine the abundance of the WT α1(I)- and G425S α1(I)-containing peptides. The difference between the WT α1(I) and G425S α1(I) peptides was quantified by analyzing the area under the curve of the liquid chromatography trace for each peptide.

### Data Availability

The RNA-Seq primary data were deposited in NCBI’s GEO repository under the accession number GSE163812.

## RESULTS

### Global Unfolded Protein Response Activation Exacerbates the OI Collagen-I Secretion Defect in Fibroblasts Expressing the *COL1A1* p.G425S Mutation

To begin, we compared the collagen-I secretion profile of wild-type (WT) α1(I) healthy donor primary fibroblasts to that of OI patient primary fibroblasts heterozygous for WT α1(I) and G425S α1(I). This common, prototypical OI-causing amino acid substitution breaks the interstrand hydrogen bond between the triple helix-forming polypeptides and causing severe forms of OI, most commonly the lethal type-II form of OI^44^. We observed that, upon ascorbate (Asc) stimulation of collagen production, collagen-I secretion over a 24 h period was significantly reduced in the heterozygous G425S α1(I)-expressing fibroblasts compared to WT α1(I)-expressing control fibroblasts (**Figure 2a**; note that in this and other media immunoblots it is not possible to normalize to a ‘housekeeping’ protein like actin due to the absence of stable markers in media – instead, we carefully ensured equivalent cell counts, media volumes, and time courses between samples as described in the Materials and Methods). Similar OI-associated collagen-I secretion defects have been observed across many OI genotypes^15, 16, 45–51^.

qPCR analysis of mRNA levels showed that classic UPR markers, including *HSPA5*, *DNAJB9*, and *HYOU1*, were not upregulated (**Figure 2b**)^23^. Thus, as expected based on prior work with other triple-helical collagen-I OI-causing variants^18, 19, 26–28^, the collagen-I secretion defect observed in G425S α1(I) OI patient fibroblasts did not induce a prototypical UPR.

We hypothesized that this absence of a prototypical UPR actually presents an opportunity for intervention. Specifically, there is still the possibility that enhancement of the ER proteostasis network via artificial UPR activation could prove beneficial for G425S α1(I) folding and trafficking. To test this hypothesis, we treated WT α1(I)-expressing or G425S α1(I)-expressing primary fibroblasts for 24 h with low concentrations of the ER stressor thapsigargin (Tg) ranging from 0.3–100 nM. Tg inhibits Ca(II) flux and acts as a rapid chemical activator of the global UPR, upregulating ER chaperones and quality control machineries^52^. We observed that even the very lowest concentration of Tg that reliably activated the UPR (10.0 nM) either had no effect or caused a strong decrease in collagen-I secretion from not just G425S α1(I) fibroblasts but also from WT α1(I) fibroblasts (**Figure 2c**; see **Figures S1a** and **S1b** for qPCR assessment of UPR activation). This decrease in collagen-I secretion was not specific to the use of Tg treatment as a strategy for global UPR activation. A second chemical UPR activator, tunicamycin (Tm; an N-glycosylation inhibitor ^53^), also reduced collagen-I secretion from both WT and OI patient primary cell lines even at the lowest concentration (0.6 μg/mL) capable of inducing the UPR (**Figure 2c**; see **Figures S1c** and **S1d** for qPCR assessment of UPR activation).

The finding that global UPR activation negatively impacted collagen-I secretion, while arguably disappointing, is unsurprising. Chemical ER stressors induce the UPR by causing general ER protein misfolding. In some cases, proteostasis defects associated with individual ER client proteins (e.g., lysosomal enzymes^35^) can still be addressed by this approach to ER proteostasis network remodeling. More commonly, however, stress-mediated induction of the entire UPR, including the PERK arm of the UPR that inhibits protein translation^22^, only exacerbates disease-causing ER proteostasis defects. Thus, the use of non-selective ER stressors was not a viable method for correcting the collagen-I secretion defect in G425S α1(I) OI patient primary fibroblasts.

### Stress-Independent Induction of the XBP1s Transcriptional Pathway Can Ameliorate the OI Collagen-I Secretion Defect

Whereas global UPR activation using chemical ER stressors is often problematic, a growing body of work suggests there is great promise in the stress-independent induction of individual UPR arms^25, 29, 31–33, 54–58^. A number of genetic and/or chemical genetic approaches are now available to force the expression of the active forms of the ATF6 and XBP1s transcription factors, the two arms of the UPR that enhance the expression of ER chaperones and quality control machineries and that are therefore most likely to result in productive ER proteostasis network remodeling^56^. These strategies have shown promise in neurodegenerative disorders^32, 57, 58^, other protein aggregation-related diseases^31, 54, 59^, and in a variety of additional contexts^33, 55^. Arm-specific UPR pathway activation has not, however, been assessed as a strategy to address collagen proteostasis defects in OI or the other collagenopathies.

We hypothesized that activation of either the ATF6 or the IRE1-XBP1s transcriptional pathways could be beneficial in G425S α1(I) OI patient fibroblasts. We first tested ATF6 activation using a chemical genetic method involving the expression in patient fibroblasts of a fusion protein composed of the constitutively active form of ATF6, ATF6(1–373)^60^, fused to a trimethoprim (TMP)-binding *Escherichia coli* DHFR-based destabilizing domain^31, 61, 62^. This approach enables TMP-dependent activation of the ATF6 transcriptional pathway (**Figure S2a**). We transduced fibroblasts with DHFR.ATF6(1–373)- or DHFR.YFP-encoding, replication-incompetent adenoviruses, and then treated with TMP. We observed selective, TMP-dependent upregulation of strongly ATF6-dependent transcriptional targets^31^, such as *HSPA5* and *HSP90B1*, with minimal induction of other UPR targets (**Figure S2b**), consistent with successful activation of the ATF6 transcriptional response. However, ATF6 activation had no detectable effect on collagen-I secretion from either WT or G425S α1(I)-expressing primary fibroblasts (**Figure S2c**), under any condition tested.

We next asked whether stress-independent induction of the IRE1-XBP1s transcriptional response might prove beneficial. For these experiments, we employed replication-incompetent adenoviruses encoding the active, spliced form of XBP1 (Ad.XBP1s)^63^ or encoding GFP (Ad.GFP) as a control. We observed that forced expression of XBP1s can often substantially increase α1(I) secretion up to ~300% from basal levels (with sensitivity to the exact Ad.XBP1s MOI used) from G425S α1(I) patient primary fibroblasts (**Figure 3a**, *Upper panels*). Notably, whenever observed, this effect was selective specifically to fibroblasts that express mutant collagen-I. α1(I) secretion by WT α1(I)-expressing primary fibroblasts was not influenced by XBP1s expression. qPCR analysis confirmed selective induction of strongly XBP1s-dependent gene transcripts upon Ad.XBP1s transduction, as opposed to global UPR activation. As shown in **Figure 3b**, transcripts such as *DNAJB9* were upregulated to levels near those attained by UPR activation using the ER stressor Tg^31^. In contrast, strongly ATF6-dependent genes, such as *HSPA5* and *HSP90B1*, or strongly ATF4-dependent genes, such as *DDIT3* and *PP1R15A*, were upregulated to a much smaller extent compared to the effects of Tg treatment ^31, 64^.

**Figure 3.**
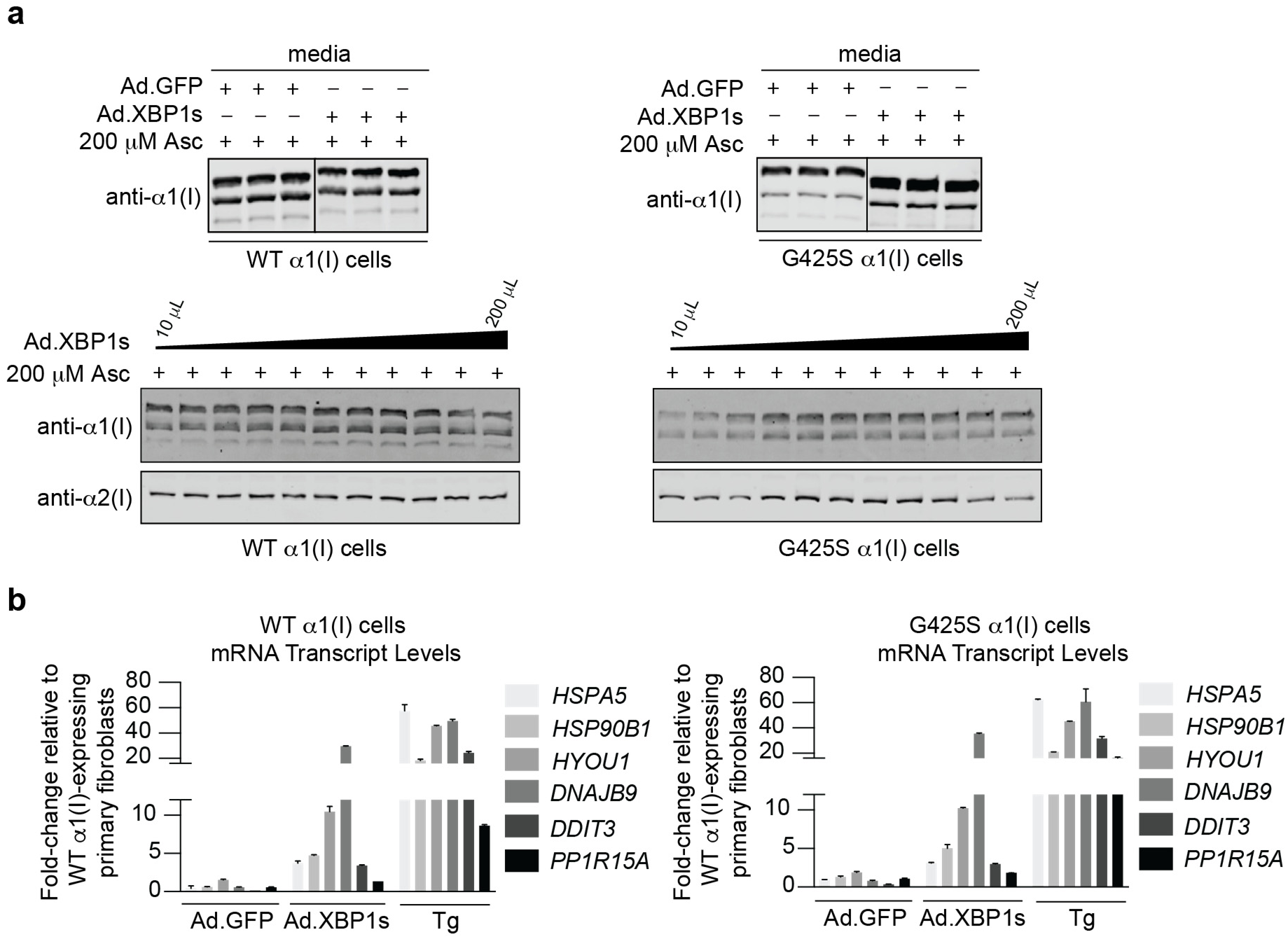
**a.** (*Upper panels*) Immunoblot showing secretion of α1(I) from WT α1(I) (*top left panel*) or G425S α1(I) (*top right panel*) primary fibroblasts transduced with Ad.GFP or Ad.XBP1s. WT α1(I) and G425S α1(I) patient fibroblasts were transduced with replication-incompetent adenoviruses engineered to deliver a constitutively expressed XBP1s gene (Ad.XBP1s) or a constitutively expressed GFP gene (Ad.GFP). Transduction with Ad.GFP was used as a negative control. α1(I) secretion was initiated by treatment of cells with sodium ascorbate (Asc) for 24 h prior to harvesting samples for analysis. The presence of multiple α1(I) bands corresponds to proα1(I) and α1(I) before and after cleavage of the C-propeptide and/or the N-propeptide. Lanes separated by a line were run on the same gel but were non-contiguous in that gel. (*Lower panels)* Dose-response immunoblots showing secretion of α1(I) and α2(I) from WT α1(I) (*bottom left panel*) and G425S α1(I) (*bottom right panel*) primary fibroblasts transduced with increasing volumes of Ad.XBP1s. Induction of collagen-I secretion was initiated by treatment of cells with ascorbate for 24 h prior to harvesting samples for analysis. **b.** qPCR analysis of UPR-dependent gene transcripts from cells treated as in the upper panels of Figure 3a. Fold-change in transcript levels was normalized to WT α1(I) fibroblasts not treated with sodium ascorbate. Thapsigargin (Tg) treatment was used as a positive control for UPR activation. Error bars represent SD across three technical replicates.

To further evaluate this phenomenon, we transduced fibroblasts with a wide range of Ad.XBP1s MOIs to control the extent of activation of the XBP1s-dependent transcriptional response. We observed a strongly Ad.XBP1s dose-dependent increase in collagen-I secretion by the G425S α1(I) primary patient fibroblasts (**Figure 3a**, *Lower right panel*; see **Figure S3** for qPCR assessment of UPR activation). In contrast, collagen-I secretion from WT α1(I) primary fibroblasts did not increase across a wide range of Ad.XBP1s doses (**Figure 3a**, *Lower left panel*). Notably, for both WT and G425S primary fibroblasts, collagen-I secretion began to drop once cells were transduced with very high doses of Ad.XBP1s. We have previously observed that tuning the level of expression of UPR transcription factors prevents any resultant cytotoxicity or global UPR activation caused by overloading the ER^31^. Consistent with this observation, Ad.XBP1s MOIs that maximize induction of strongly XBP1s-dependent genes, such as *DNAJB9*, while minimizing upregulation of genes that are also strongly regulated by ATF6, such as *HSPA5* (see **Figure S3**), often resulted in optimal improvements of collagen-I secretion by G425S α1(I) primary patient fibroblasts.

We note that further studies in the process of preparing this publication for review revealed unresolved inconsistency in the XBP1s-induced enhancement of collagen-I secretion from G425S α1(I) primary patient fibroblasts. While the result shown in **Figure 3A** was replicable by different researchers and laboratories, at times those same researchers would not observe the XBP1s-enhanced secretion. After extensive optimization and testing of various experimental variables, including MOI for viral transduction, age and aliquots of the primary fibroblasts used, plating density, timecourses, and more, we have not been able to identify a single variable that explains why the **Figure 3A** result is sometimes observed and sometimes is not. The mechanistic investigation presented below should be considered with this caveat in mind.

### XBP1s Pathway Activation May Be Able to Restore G425S α1(I) Trafficking by Selectively Enhancing Mutant Collagen-I Folding

We sought to gain initial insights into the underlying mechanism(s) that can drive(s) an XBP1s-dependent selective increase in collagen-I secretion only from G425S α1(I) OI-patient primary fibroblasts, but not from WT α1(I) primary fibroblasts. We first used RNA-Seq to evaluate whether the transcriptional consequences of Ad.XBP1s treatment were substantively different in WT α1(I)-expressing versus G425S α1(I)-expressing primary fibroblasts (**Table S1**). Correlation analysis revealed that the transcriptional remodeling caused by Ad.XBP1s was broadly similar across these two cell lines (**Figure 4a**). Moreover, gene set enrichment analysis^40^ confirmed that Ad.XBP1s treatment upregulated expected ER proteostasis-related gene sets, with strongly enriched gene sets shared across both samples and the top five enriched gene sets being identical (**Table S2**; see **Figure S4** for images of relevant enrichment plots). Thus, it seems unlikely that observations of a selective increase in collagen-I secretion only from the G425S α1(I) OI patient fibroblasts can be attributed to broadly different transcriptional consequences of XBP1s activation in those cells.

**Figure 4.**
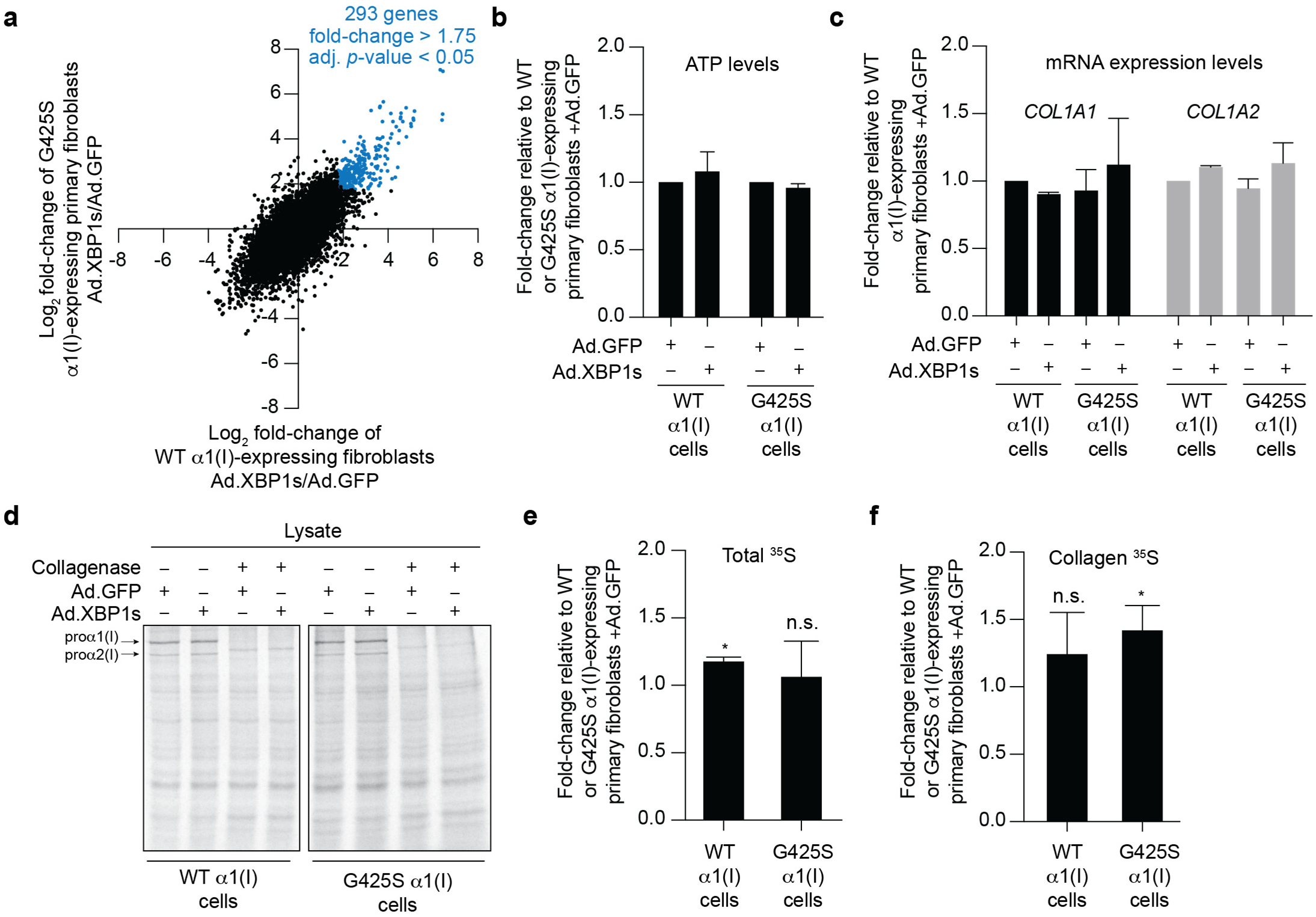
**a.** Correlation plot showing all transcripts expressed in sodium ascorbate-treated WT α1(I) or G425S α1(I) primary fibroblasts transduced with Ad.XBP1s versus Ad.GFP. Genes that were significantly upregulated (>1.75-fold and Benjamin-Hochberg adjusted *p*-value <0.05) in both WT α1(I) and G425S α1(I) upon treatment with Ad.XBP1s vs Ad.GFP are shown in blue. **b.** Quantification of cellular ATP using the CellTiter-Glo® assay for sodium ascorbate-treated WT α1(I) or G425S α1(I) primary fibroblasts transduced with Ad.XBP1s versus Ad.GFP. Fold-change was normalized to Ad.GFP transductions. Error bars represent SD across three biological replicates. **c.** qPCR analysis of *COL1A1* and *COL1A2* gene transcripts in sodium ascorbate-treated WT α1(I) or G425S α1(I) primary fibroblasts transduced with Ad.XBP1s versus Ad.GFP. Error bars represent SD across three biological replicates. Fold-change was normalized to WT α1(I) cells treated with Ad.GFP. **d.** Representative SDS-PAGE gel of ^35^S-Cys/Met-labeled cell lysates from sodium ascorbate-treated WT α1(I) or G425S α1(I) primary fibroblasts transduced with Ad.XBP1s versus Ad.GFP. Samples were split in half and treated with or without bacterial collagenase for three days to determine which protein bands derived from collagen-I, with the resulting proα1(I) and proα2(I) bands highlighted by arrows. **e.** Quantification of the total ^35^S signal in the absence of collagenase for the samples in Figure 4d. Signal was normalized to cells treated with Ad.GFP for each cell line. Error bars represent SD over three biological replicates. * and n.s. correspond to *p*-value < 0.05 and not significant, respectively. **f.** Quantification of the collagen-I signal for the samples in Figure 4d. Signal was normalized to cells treated with Ad.GFP for each cell line. Error bars represent SD over three biological replicates. * and n.s. correspond to *p*-value < 0.05 and not significant, respectively.

We next asked whether XBP1s might be selectively improving G425S α1(I) trafficking by influencing cell growth, collagen-I transcription, global protein translation, or collagen-I translation specifically in the primary OI patient fibroblasts but not in the healthy donor control fibroblasts. Quantification of cellular ATP using a CellTiter-Glo® assay showed that Ad.XBP1s treatment had no detectable influence on cell growth rates in either G425S or WT α1(I) primary fibroblasts (**Figure 4b**). Targeted qPCR analysis of collagen-I transcripts indicated that Ad.XBP1s treatment also did not substantially alter *COL1A1* or *COL1A2* mRNA levels in either cell line (**Figure 4c**). Finally, we used ^35^S-Cys/Met metabolic labeling experiments to evaluate possible effects on protein translation that might have occurred only in G425S α1(I) primary patient fibroblasts. After labeling all nascent chains synthesized in a 10-minute span using ^35^S-Cys/Met treatment, we immediately lysed the cells and separated the proteins using SDS-PAGE (**Figure 4d**). Quantification of the resulting autoradiographs revealed that global protein synthesis was significantly but insubstantially increased in WT α1(I) fibroblasts upon treatment with Ad.XBP1s, while protein synthesis in G425S α1(I) fibroblasts remained unchanged (**Figure 4e**). Collagenase digestion of the cell lysates allowed us to identify the procollagen-I bands on these SDS-PAGE gels, as indicated by the arrows in **Figure 4d**. Specifically quantifying the procollagen-I bands indicated that Ad.XBP1s treatment very modestly upregulated collagen-I synthesis in G425S α1(I) and possibly in WT α1(I) fibroblasts (**Figure 4f**).

Based on these observations, the observed ability of Ad.XBP1s treatment to, at least in some conditions, selectively enhance collagen-I trafficking up to ~300% in G425S α1(I) primary fibroblasts but not in WT α1(I) primary fibroblasts cannot likely be attributed to differential consequences of Ad.XBP1s for the overall cellular transcriptome, cell growth, collagen-I transcription, or global protein translation. Given the effects observed, we would not completely rule out the possibility that modulation of collagen-I synthesis by Ad.XBP1s is involved. It appears more likely, however, that XBP1s-mediated remodeling of the ER proteostasis network selectively modulates downstream steps in collagen-I folding, quality control, or trafficking that are defective in the G425S α1(I)-producing primary patient fibroblasts.

### XBP1s Activation Can Increase the Ratio of G425S Mutant to WT α1(I) Polypeptides Secreted in the Form of Stable Collagen Triple Helices

To further improve our understanding of how Ad.XBP1s selectively promotes collagen-I secretion from G425S α1(I)-expressing but not WT α1(I)-expressing primary fibroblasts, we sought to assess whether Ad.XBP1s alters the stability or composition of secreted collagen-I. This clinically severe form of OI is an autosomal dominant disease, and G425S patient fibroblasts synthesize both WT and mutant α1(I). If Ad.XBP1s treatment does indeed enhance the folding or assembly of G425S α1(I) into triple-helical collagen, we would predict that the enhanced collagen-I secretion often observed would result in a higher ratio of G425S:WT α1(I) in the secreted collagen-I obtained from Ad.XBP1s-transduced primary fibroblasts relative to Ad.GFP-transduced fibroblasts.

To first evaluate whether Ad.XBP1s enhances the secretion of properly folded triple-helical collagen-I versus badly misfolded and/or non-triple-helical collagen-I from G425S α1(I)-expressing primary patient fibroblasts, we turned to proteolysis assays. One characteristic of the fibrillar collagens is that stable, folded triple helices are resistant to protease digestion^42, 65^. In contrast, denatured or badly misfolded collagen-I is susceptible to cleavage by pepsin and trypsin/chymotrypsin. We performed two separate sets of assays to assess the consequences of Ad.XBP1s treatment for the triple-helical structure of procollagen-I secreted from G425S α1(I) primary patient fibroblasts. In the first assay, procollagen-I precipitated from media fractions was treated with trypsin or chymotrypsin and the extent of degradation was analyzed by immunoblotting with detection using a polyclonal antibody raised against the triple-helical domain of α2(I). We found that, in the context of either Ad.GFP or Ad.XBP1s transduction, procollagen-I samples harvested from WT and G425S α1(I)-producing primary fibroblasts were stable to prolonged treatment with trypsin and chymotrypsin to similar extents (**Figure 5a**; note that trypsin and chymotrypsin cleave the collagen-I propeptide domains causing the observed drop in molecular weight), consistent with Ad.XBP1s having no negative effects on the extent to which collagen-I attained a stable, triple-helical form.

**Figure 5.**
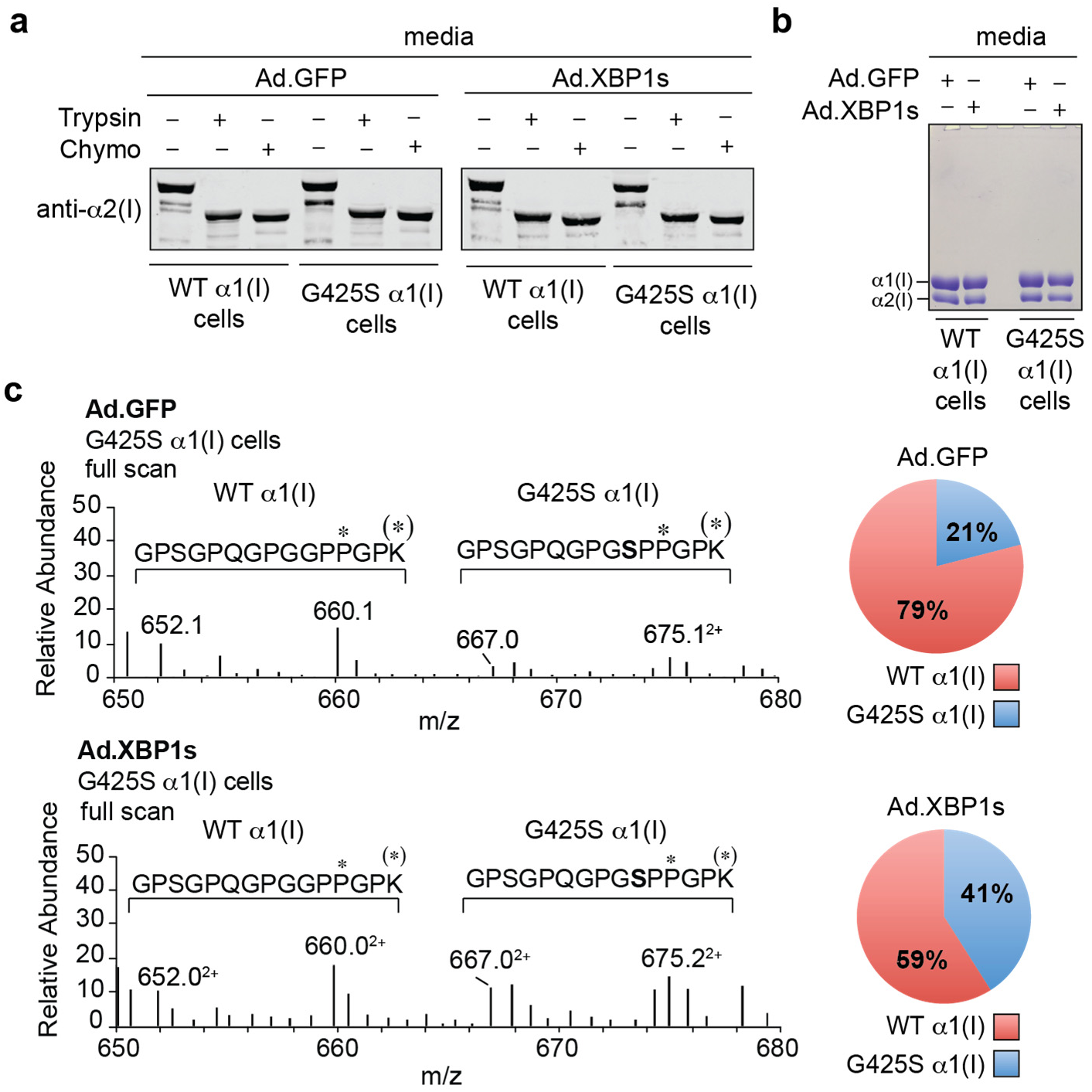
**a.** Immunoblot showing protease digestion of secreted α1(I) and α2(I) from WT α1(I) or G425S α1(I) primary fibroblasts transduced with Ad.GFP or Ad.XBP1s. Collagen-I was precipitated from the media and then treated with trypsin or chymotrypsin for 2 h at room temperature, following which the reaction was quenched prior to analysis by immunoblotting using a polyclonal antibody against the triple-helical domain of α2(I). **b.** Coomassie-stained SDS-PAGE gel showing pepsin-digested collagen-I secreted from sodium ascorbate (Asc)-treated WT α1(I) or G425S α1(I) primary fibroblasts transduced with Ad.GFP or Ad.XBP1s. Collagen-I was precipitated from media, resolubilized, and treated with pepsin. Note that α1(I) is known to display lower electrophoretic mobility than α2(I)^88^. **c.** Representative MS1 scan highlighting the four relevant *m*/*z* ratios and quantification of the WT α1(I) and G425S α1(I) allelic products secreted from G425S α1(I) primary patient fibroblasts transduced with Ad.GFP or Ad.XBP1s. The α1(I) band from Figure 5b was excised, reduced, alkylated, and trypsinized. The resultant peptides were analyzed by LC-MS/MS analysis to evaluate the abundance of the WT G425 versus mutant S425 α1(I)-containing peptides. *m*/*z* ratios of 652.1 and 667.0 correspond to the peptide without lysine hydroxylation, while the ratios of 660.1 and 675.1 accounts for the hydroxylated lysine. The difference between the WT α1(I) and G425S α1(I) peptides was quantified by analysis of the area under the curve of the LC trace for each peptide.

In the second assay, we sought to assess whether and how Ad.XBP1s treatment altered the extent to which G425S α1(I) was incorporated into the secreted collagen-I triple helices. Briefly, media from G425S α1(I)-expressing primary fibroblasts transduced with either Ad.GFP as a control or Ad.XBP1s was collected. Samples were treated with pepsin to degrade non-collagenous proteins and any non-triple-helical collagen, and then the secreted triple-helical collagen-I was precipitated and resolubilized. Separation of the digested protein samples using SDS-PAGE revealed the presence of pepsin-stable α1(I) and α2(I) polypeptides isolated by this protocol (**Figure 5b**), further confirming that Ad.XBP1s treatment does not result in excessive production of badly misfolded or unfolded collagen-I from G425S α1(I) primary patient fibroblasts. We next applied a well-established MS protocol^66–69^ to evaluate whether XBP1s alters the ratio of G425S α1(I):WT α1(I) secreted in a pepsin-stable, triple-helical form. LC-MS/MS analysis of the composition of the α1(I) band from **Figure 5b** showed that, in media samples obtained from Ad.GFP-treated G425S α1(I) primary fibroblasts, the G425S allelic product constituted roughly 20% of the total α1(I) produced. Ad.XBP1s treatment doubled the fraction of the G425S allelic product present, shifting the ratio of G425S:WT α1(I) to nearly 1:1 (**Figure 5c**). This increase in secretion of the G425S α1(I) allelic product confirms that Ad.XBP1s treatment specifically can promote the folding and/or trafficking of the mutant collagen-I produced by the G425S α1(I) OI patient primary fibroblasts.

## DISCUSSION

The G425S α1(I) variant causes a severe, autosomal dominant form of OI^44^. The detailed origin of pathology is unclear, and may involve both intracellular and extracellular defects. Like many other OI-causing variants^15, 16, 45–49^, the G425S amino acid substitution results in a substantial collagen-I folding/secretion defect^51^. Such a proteostasis defect might also be expected to lead to cellular dysfunction, reducing overall bone health and motivating an effort to discover strategies to improve collagen-I trafficking in this genotype.

We find that collagen-I secretion by G425S α1(I)-expressing primary patient fibroblasts can, under some experimental conditions,be rescued by UPR arm-specific, ER stress-independent induction of the XBP1s transcriptional response. This effect is unique to XBP1s, as neither induction of the global UPR using small molecule stressors nor arm-selective, stress-independent induction of the ATF6 transcriptional pathway of the UPR has any beneficial effect. While XBP1s can enhance quality control for a number of aggregation-prone ER client proteins ^31, 34, 57, 58^, to our knowledge this observation constitutes the first example of XBP1s-mediated ER proteostasis network remodeling demonstrating an ability to resolve a specific protein secretion defect.

Despite similar transcriptional effects in both G425S α1(I)-expressing primary patient fibroblasts and healthy donor primary fibroblasts, the consequences of XBP1s activity are, whenever observed, highly selective for mutant collagen-I. That is, only the OI-patient fibroblasts displayed an XBP1s-dependent increase in collagen-I secretion. Collagen-I secretion from WT fibroblasts was consistently unaltered by XBP1s treatment. The effect appears most likely to be post-translational, perhaps involving improved chaperone-mediated folding and assembly of the mutant collagen-I. This notion is consistent with our analysis of the composition of the triple-helical collagen-I secreted upon XBP1s treatment, in which secretion of the mutant G425S α1(I) allele is selectively enhanced relative to the WT α1(I) allele.

Despite decades of research, current therapeutic strategies to address OI still have many deficits^70, 71^. The most common clinical approach involves treatment with anti-resorptive agents, most prominently the bisphosphonates^71^. In contrast to inhibitors of catabolism, inducers of bone anabolism have recently received increasing attention. Of particular significance, increasing bone production via enhancing LRP5 signaling genetically or by targeting sclerostin and using antibodies to inhibit TGFβ signaling have both shown promise, with the TGFβ-targeting strategy currently in clinical trials^54, 70, 72, 73^. Prior work has also suggested the potential of the chemical chaperone 4-phenylbutyric acid to ameliorate skeletal defects in an OI zebrafish model^74, 75^.

The data presented here suggest that ER stress-independent IRE1-XBP1s activation may offer a new anabolism-targeted strategy to consider – one predicated on a beneficial remodeling of the ER proteostasis network to promote collagen-I folding and secretion, restore cell health, and potentially promote bone production. There is much work to be done following this initial *in vitro* proof-of-principle. Although primary fibroblasts are a highly relevant cell type that natively produces large quantities of collagen-I, and even though skin defects are commonly associated with OI^3, 76^, it will be important to evaluate the consequences of XBP1s activity in bone-producing osteoblasts. Moreover, unidentified experimental variables seem to determine whether the effect is observed in any given experiment, and we have so far been unable to resolve this issue.Primary osteoblasts are not widely available for this or many other OI variants. New OI animal models that incorporate prototypical OI-causing collagen-I triple-helix mutations, such as Gly→Ser-inducing mutations in *COL1A1*, would be particularly beneficial in this regard and are currently lacking. The only triple helix-domain single amino acid substitution-based mouse models for OI that have been reported to date are the G349C α1(I) and the G610C α2(I) mice^77, 78^. With the recent emergence of methods to grow bone-like matrices in a dish, perhaps an opportunity exists to screen XBP1s activators in a more experimentally reproducible OI-mutant environment^79^. It will be important to examine the scope of OI mutants for which XBP1s activation is beneficial. As with other strategies for OI amelioration, it is likely that the approach is applicable to only a subset of variants. Progress will also benefit from improved small molecule-based IRE1-XBP1s activators, as current best-in-class compounds provide only short-term induction of the pathway and it is currently unclear whether they would function in osteoblasts^80^. Molecular-level mechanistic understanding of XBP1s-enhanced assembly and secretion would also be valuable, and will benefit from an improved understanding of fundamental collagen secretion and quality control mechanisms^81–86^. Such understanding could allow an even more specific strategy to resolve collagen proteostasis defects that does not rely upon large-scale XBP1s-mediated remodeling of the ER proteostasis network.

In conclusion, we have observed that XBP1s-mediated enhancement of the ER proteostasis network can, at least under some experimental conditions, selectively improve the folding and secretion of an OI-causing collagen-I variant in primary patient fibroblasts. Considering the vast array of mutations that engender collagen proteostasis defects and cause currently incurable collagenopathies, this observation should catalyze further efforts to understand and test the broad potential of proteostasis network modulation in these disorders.

## ACKNOWLEDGEMENTS

This work was supported by the National Institutes of Health (Grant 1R01AR071443), a Research Grant from the G. Harold and Leila Y. Mathers Foundation, a Dreyfus Foundation Teacher-Scholar Award, and an MIT International Science and Technology Initiatives grant (all to M.D.S.), and by the National Health and Medical Research Council of Australia (Grant GNT 1144807 to J.F.B. and S.R.L.) and the Victorian Government’s Operational Infrastructure Support Program. A.S.D. was supported by a NIH Ruth L. Kirschstein predoctoral fellowship (F31AR067615). N.-D.D. was supported by Canadian Institutes of Health Research and Fonds de Recherche du Québec–Santé postdoctoral fellowships. A.A.B. was supported by the Johnson and Johnson UROP Scholar Program. Additional support was provided by an NIEHS grant (P30-ES002109) to the MIT Center for Environmental Health Sciences.

