## Supplemental Information for "XBP1s-Mediated ER Proteostasis Network Enhancement Can Selectively Improve the Folding and Secretion of an Osteogenesis Imperfecta-Causing Collagen-I Variant"


| **Page** | **Contents** |
|---|---|
| S1 | Table of Contents |
| S2 | Supporting Tables |
| S3–7 | Supporting Figures |

**Table S1.** RNA-Seq differential gene expression analysis for sodium ascorbate-treated WT $\alpha$1(I)- or G425S $\alpha$1(I)-expressing primary fibroblasts transduced with Ad.XBP1s versus Ad.GFP (see attached Excel file).

**Table S2.** RNA-Seq gene set enrichment analysis for samples in **Table S1** (see attached Excel file).

**Table S3.** qPCR primer sequences.

| Gene | Forward | Reverse |
|---|---|---|
| *RPLP2* | 5′-CCATTCAGCTCACTGATAACCTT-3′ | 5′-CGTCGCCTCCTACCTGCT -3′ |
| *HSP90B1* | 5′-GGCCAGTTTGGTGTCGGT -3′ | 5′-CGTTCCCCGTCCTAGAGTGTT -3′ |
| *HSPA5* | 5′-GCCTGTATTTCTAGACCTGCC -3′ | 5′-TTCATCTTGCCAGCCAGTTG-3′ |
| *DNAJB9* | 5′-CTGTATGCTGATTGGTAGAGTCAA -3′ | 5′-AGTAGACAAAGGCATCATTTCCAA-3′ |
| *SEC24D* | 5′-AGCAGACTGTCCTGGGAAGC -3′ | 5′-TTTGTTTGGGGCTGGAAAAG-3′ |
| *DDIT3* | 5′-GGAGCTGGAAGCCTGGTATG -3′ | 5′-GCCAGAGAAGCAGGGTCAAG-3′ |
| *HYOU1* | 5′-GCAGACCTGTTGGCACTGAG -3′ | 5′-TCACGATCACCGGTGTTTTC-3′ |
| *PPP1R15A* | 5′-CTGAAGCCTGGGGACTTTTG -3′ | 5′-CTTTCTCCTCCCCTGGGTTC-3′ |
| *COL1A1* | 5′-TGGTAGCCGTGGTTTCCCTG -3′ | 5′-TCCAGTCAGACCCTTGGCAC-3′ |
| *COL1A2* | 5′-TGGCTCGAGAGGTGAACGTG-3′ | 5′-AGCACCGTTGACTCCAGGAC-3′ |

**Figure S1**
**a–b.** qPCR analysis of unfolded protein response (UPR)-dependent genes in WT (**Figure S1a**) or G425S (**Figure S1b**) α1(I)-expressing primary fibroblasts treated with increasing concentrations of thapsigargin (Tg). Transcript levels were normalized to vehicle-treated WT α1(I)-expressing cells not treated with sodium ascorbate (Asc). Error bars represent SD across three technical replicates.
**c–d.** qPCR analysis of unfolded protein response (UPR)-dependent genes in WT (**Figure S1c**) or G425S (**Figure S1d**) α1(I)-expressing primary fibroblasts treated with increasing concentrations of tunicamycin (Tm). Transcript levels were normalized to vehicle-treated WT α1(I)-expressing cells not treated with Asc. Error bars represent SD across three technical replicates.

**Figure S2**
**a.** Illustration of trimethoprim (TMP)-regulated DHFR.ATF6(1–373) constructs. In the absence of TMP, DHFR.ATF6 is rapidly degraded by the proteasome. TMP treatment stabilizes the folded form of DHFR, allowing the DHFR.ATF6 construct to persist and perform its transcription factor function.
**b.** qPCR analysis of unfolded protein response (UPR)-regulated transcripts in WT α1(I) (left) or G425S α1(I) (right) primary fibroblasts transduced with a replication-incompetent adenovirus (Ad) encoding either DHFR.YFP (as a control) or DHFR.ATF6(1–373). 24 h post-transduction, media was replaced with fresh media containing 10 μM trimethoprim (TMP). 48 h post-media change, media was removed and replaced with fresh media containing 200 μM sodium ascorbate (Asc). Samples were harvested the next day for qPCR analysis. Error bars represent SD across three technical replicates
**c.** Immunoblot showing secretion of α1(I) and α2(I) from WT α1(I)- or G425S α1(I)-expressing primary fibroblasts transduced with either Ad.DHFR.YFP or Ad.DHFR.ATF6(1–373). 48 h post-media change, media was removed and replaced with fresh media containing 200 μM sodium ascorbate. Samples were harvested the next day for qPCR analysis. The presence of multiple α1(I) bands likely corresponds to proα1(I) and α1(I) following cleavage of the C-propeptide and/or the N-propeptide.

**Figure S3**
qPCR analysis of unfolded protein response (UPR)-regulated gene transcripts in G425S α1(I) primary patient fibroblast cells treated with increasing volumes of Ad.GFP (**Left**) or Ad.XBP1s (**Right**). Fold-change was normalized to untransduced G425S α1(I) cells not treated with sodium ascorbate (Asc). Error bars represent SD across three technical replicates.

**Figure S4**
Gene set enrichment plots for the top five XBP1s activation-related gene sets in WT α1(I) and G425S α1(I) primary fibroblasts. Induction of XBP1s expression activates expected ER proteostasis-related gene sets. These gene sets are drawn from the MSigDB C5 collection. See **Table S2** for the complete gene set enrichment analysis.

**Data availability**
Supplementary information is available at the Experimental & Molecular Medicine website. Supplementary information accompanies the manuscript on the Experimental & Molecular Medicine website http://www.nature.com/emm/.

**Figure S1**

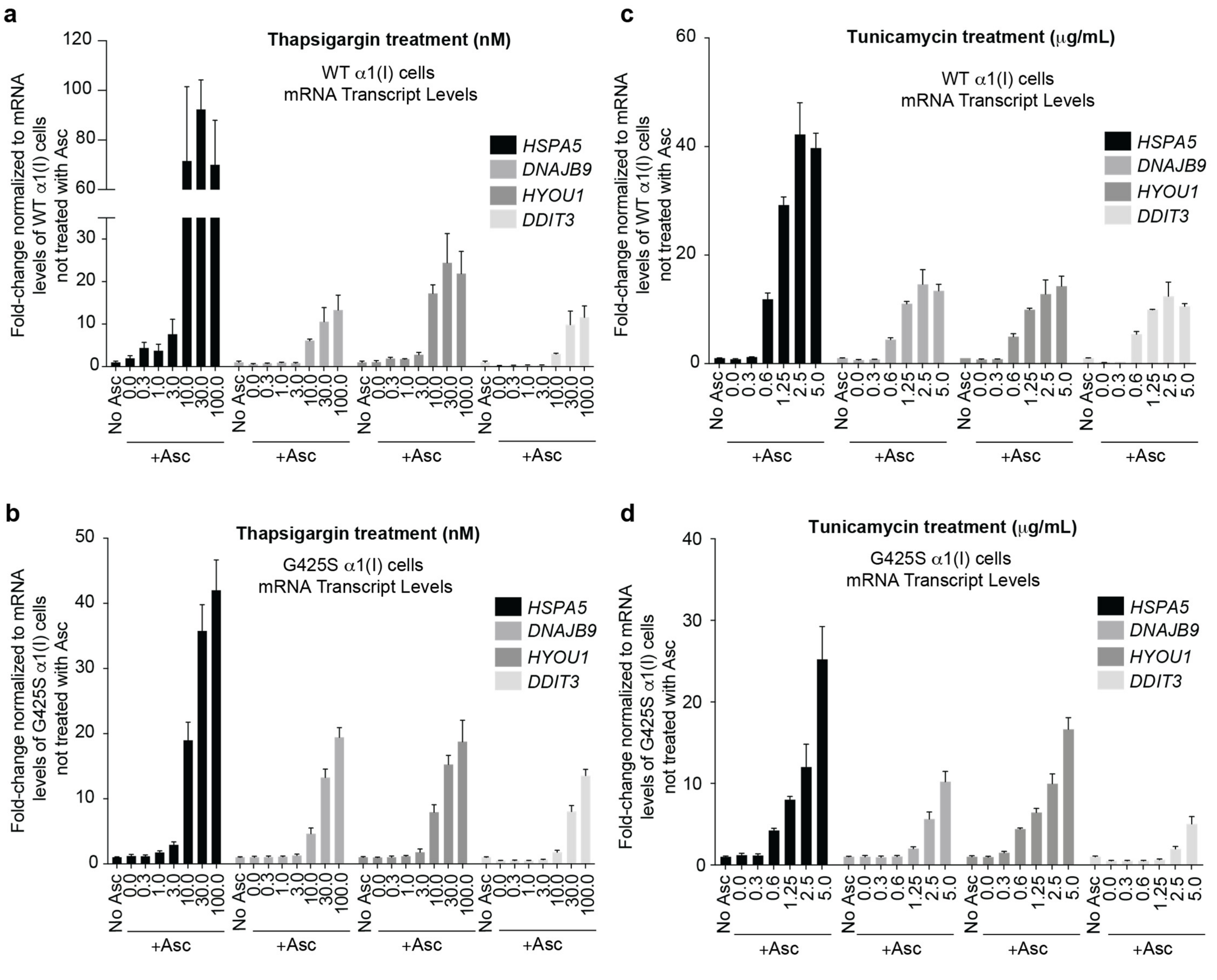
a
Thapsigargin treatment (nM)
WT α1(I) cells
mRNA Transcript Levels
Fold-change normalized to mRNA levels of WT α1(I) cells not treated with Asc
HSPA5
DNAJB9
HYOU1
DDIT3
No Asc
+Asc
b
Thapsigargin treatment (nM)
G425S α1(I) cells
mRNA Transcript Levels
Fold-change normalized to mRNA levels of G425S α1(I) cells not treated with Asc
c
Tunicamycin treatment (μg/mL)
WT α1(I) cells
mRNA Transcript Levels
Fold-change normalized to mRNA levels of WT α1(I) cells not treated with Asc
d
Tunicamycin treatment (μg/mL)
G425S α1(I) cells
mRNA Transcript Levels
Fold-change normalized to mRNA levels of G425S α1(I) cells not treated with Asc

**Figure S2**

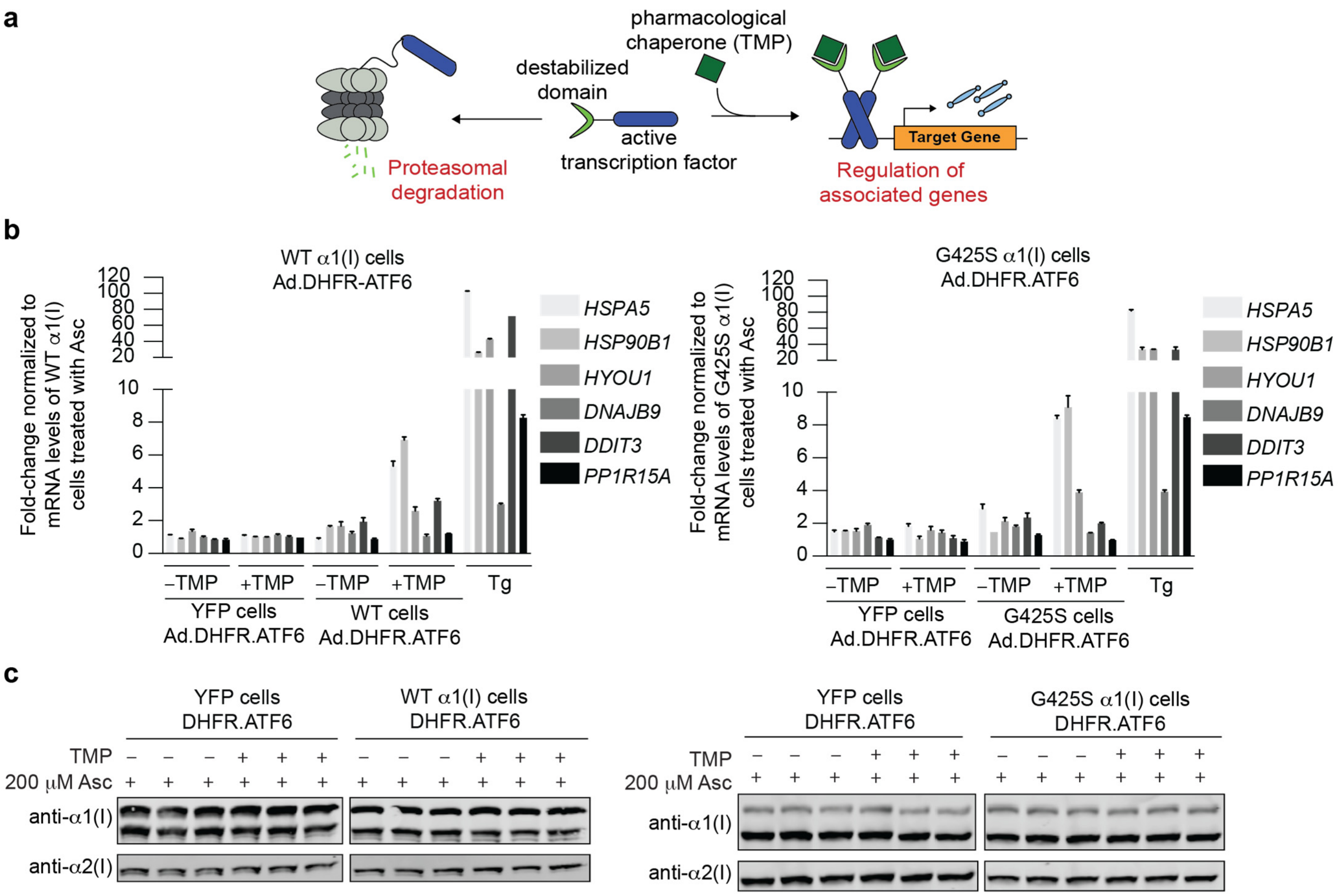
a
pharmacological chaperone (TMP)
destabilized domain
active transcription factor
Proteasomal degradation
Target Gene
Regulation of associated genes
b
WT α1(I) cells Ad.DHFR-ATF6
Fold-change normalized to mRNA levels of WT α1(I) cells treated with Asc
HSPA5
HSP90B1
HYOU1
DNAJB9
DDIT3
PP1R15A
–TMP
+TMP
YFP cells Ad.DHFR.ATF6
WT cells Ad.DHFR.ATF6
Tg
G425S α1(I) cells Ad.DHFR.ATF6
Fold-change normalized to mRNA levels of G425S α1(I) cells treated with Asc
G425S cells Ad.DHFR.ATF6
c
YFP cells DHFR.ATF6
WT α1(I) cells DHFR.ATF6
G425S α1(I) cells DHFR.ATF6
TMP
200 μM Asc
anti-α1(I)
anti-α2(I)

**Figure S3**

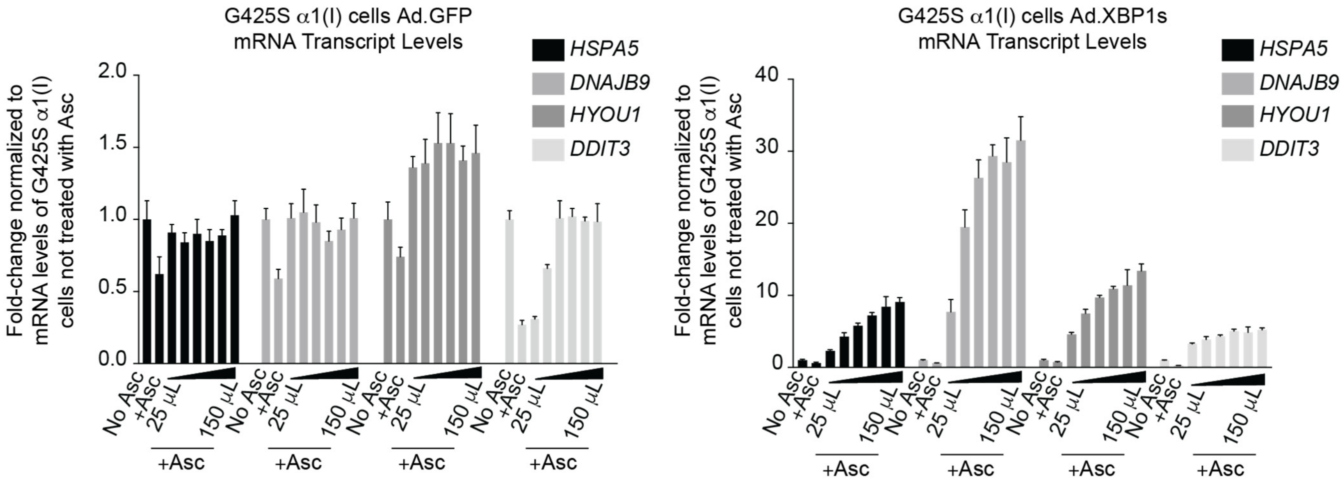
G425S α1(I) cells Ad.GFP
mRNA Transcript Levels
Fold-change normalized to mRNA levels of G425S α1(I) cells not treated with Asc
HSPA5
DNAJB9
HYOU1
DDIT3
2.0
1.5
1.0
0.5
0.0
No Asc
+Asc
25 μL
150 μL
+Asc
G425S α1(I) cells Ad.XBP1s
mRNA Transcript Levels
40
30
20
10
0

**Figure S4**

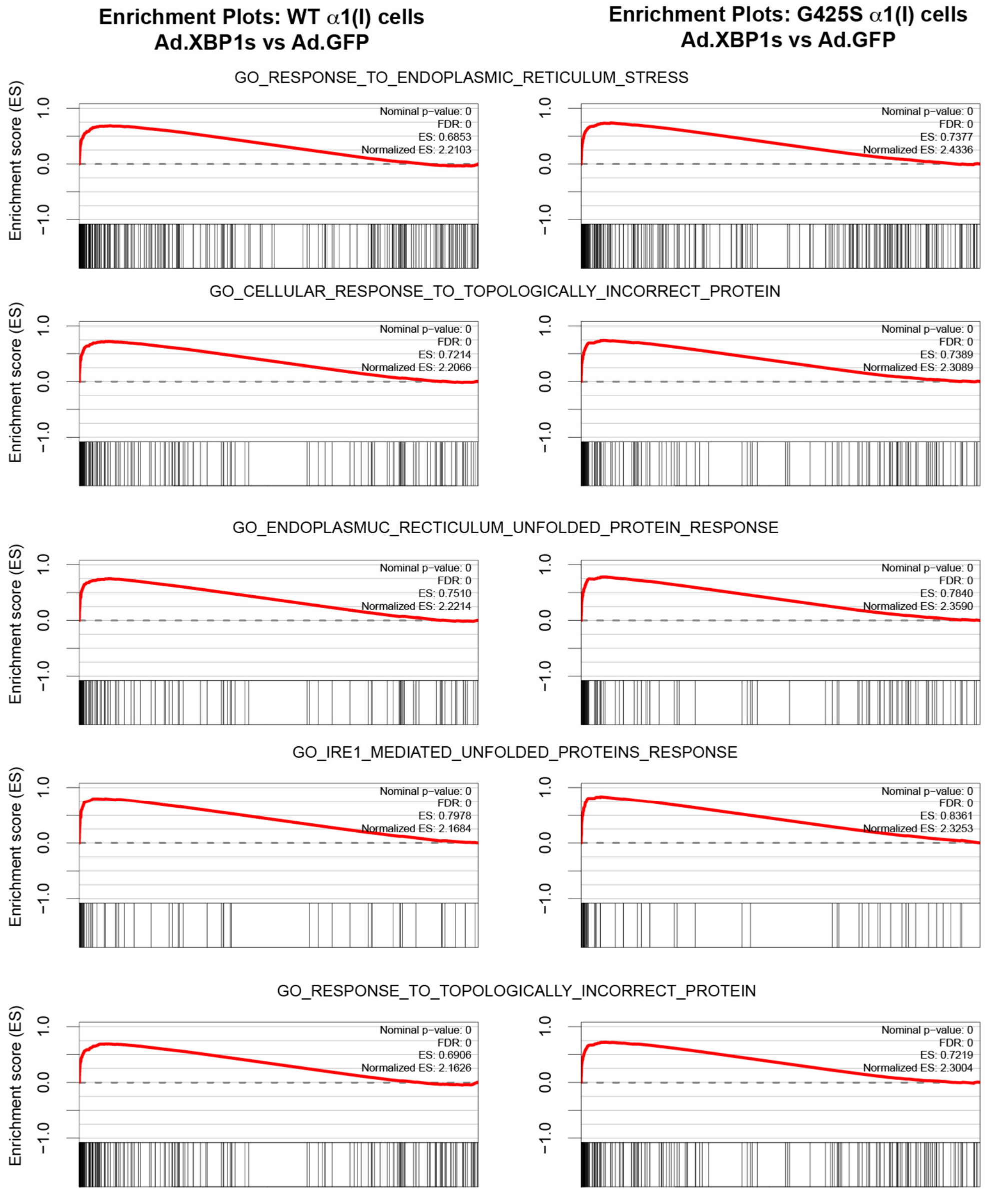
Enrichment Plots: WT α1(I) cells
Ad.XBP1s vs Ad.GFP
Enrichment Plots: G425S α1(I) cells
Ad.XBP1s vs Ad.GFP
GO_RESPONSE_TO_ENDOPLASMIC_RETICULUM_STRESS
Nominal p−value: 0
FDR: 0
ES: 0.6853
Normalized ES: 2.2103
Nominal p−value: 0
FDR: 0
ES: 0.7377
Normalized ES: 2.4336
GO_CELLULAR_RESPONSE_TO_TOPOLOGICALLY_INCORRECT_PROTEIN
Nominal p−value: 0
FDR: 0
ES: 0.7214
Normalized ES: 2.2066
Nominal p−value: 0
FDR: 0
ES: 0.7389
Normalized ES: 2.3089
GO_ENDOPLASMUC_RECTICULUM_UNFOLDED_PROTEIN_RESPONSE
Nominal p−value: 0
FDR: 0
ES: 0.7510
Normalized ES: 2.2214
Nominal p−value: 0
FDR: 0
ES: 0.7840
Normalized ES: 2.3590
GO_IRE1_MEDIATED_UNFOLDED_PROTEINS_RESPONSE
Nominal p−value: 0
FDR: 0
ES: 0.7978
Normalized ES: 2.1684
Nominal p−value: 0
FDR: 0
ES: 0.8361
Normalized ES: 2.3253
GO_RESPONSE_TO_TOPOLOGICALLY_INCORRECT_PROTEIN
Nominal p−value: 0
FDR: 0
ES: 0.6906
Normalized ES: 2.1626
Nominal p−value: 0
FDR: 0
ES: 0.7219
Normalized ES: 2.3004
Enrichment score (ES)
1.0
0.0
−1.0
